# Macrophage Signature-Based Prediction of Cancer Treatment Response Using Attention-Weighted Multiple Instance Learning

**DOI:** 10.64898/2026.08.28.747791

**Authors:** Matthew Madgwick, Samuel Witham, Martina Occhetta, Moritz Haneklaus, Barbara Camanzi, Michails Smyrnakis, Laura-Jayne Gardiner

**Affiliations:** IBM Research, Hartree Centre, Daresbury; Macomics Ltd, Technopark, Cambridge; STFC, Hartree Centre, Daresbury

**Keywords:** Foundation Model, Gene Regulatory Networks, Multiple Instance Learning, Contrastive Learning, Cancer, Macrophages

## Abstract

Predicting immunotherapy response from single-cell data remains difficult due to patient-level labels, extreme class imbalance, and highly heterogeneous macrophage states. We present a Multiple Instance Learning (MIL) framework that treats each patient as a bag of macrophage embeddings derived from a single-cell RNA foundation model. The architecture incorporates an attention-based pooling mechanism with reduced model complexity, dropout-enhanced regularization and explicit attention penalties to improve stability in small-sample regimes. To address imbalanced clinical datasets, MIL outputs are optimized with a combined focal loss and supervised contrastive objective that simultaneously sharpens class boundaries and improves representation clustering. Across three cancer datasets, this approach outperforms pseudobulk aggregation, embedding baselines and standard MIL variants. Attention-weighted attribution and transcriptional regulatory analysis reveal distinct macrophage programs, interferon and antigen-presentation networks in responders versus hypoxia-linked regulatory modules in non-responders. This shows the potential of MIL to uncover predictive and mechanistically interpretable immune states.

## 1 Introduction

Macrophages are professional phagocytic cells of the innate immune system. They reside in every tissue, where they clear debris, remodel the extracellular matrix to direct tissue repair, and coordinate the resolution of inflammation in order to preserve homeostasis [Gautier et al., 2012, Hume, 2008, Gordon and Taylor, 2005]. They are derived from peripheral monocytes or embryonic-derived tissue-resident lineages, giving rise to diverse macrophage populations across tissues [Laviron and Boissonnas]. Macrophages are abundant in the tumor microenvironment, where they can either promote or inhibit tumor growth depending on their transcriptional state [Zhang et al.]. However, in the majority of cases, a high tumor associated macrophage (TAM) burden correlates with aggressive disease and inferior survival [Qian and Pollard, 2010, Wynn et al., 2013, Stultz and Fong, 2021, Erlandsson et al., 2019, Lundholm et al., 2015, Lanciotti et al., 2014].

TAM state is influenced by the tumor itself. In early tumor development, pro-inflammatory signals polarize TAMs toward an M1-like phenotype, releasing IFN-*γ*, TNF-*α*, IL-6 and other cytokines that maintain the chronic, low-grade inflammation permissive for tumor initiation [Balkwill and Mantovani, 2012, Wynn et al., 2013]. Once the tumor is established, tumor-derived signals such as TGF-*β*, CSF-1, chemokines (e.g. CCL2), cytokines (e.g. IL-1 and IL-4), immune complexes, and complement, ‘educate’ TAMs towards an M2-like immunosuppressive phenotype that promotes tumor growth [Qian and Pollard, 2010, Mantovani et al., 2022], in part via programmed death-ligand 1 (PD-L1) and arginase-1 upregulation [Xu et al.]. In this state, TAMs enhance angiogenesis and tumor invasiveness by sustaining chronic inflammation, supporting epithelial-mesenchymal transition (EMT) and invasion, and suppressing anti-tumor immunity through expression of immunosuppressive cytokines [Prenen and Mazzone, Yang et al., Quail and Joyce]. TAMs also interact with other immunosuppressive cells, such as regulatory T cells, to create a microenvironment that fosters immune suppression [Stultz and Fong, 2021, Sun et al., 2023]. Given these largely pro-tumor functions, various immunotherapy strategies are being explored to reprogram or deplete TAMs, such as CSF1R inhibitors [Dou et al.], chimeric antigen receptor (CAR)-macrophages [Nadella and Sharma] and CD47/SIRP*α* blockers [Bouwstra et al.].

The polarization axis is a simplification: TAMs occupy a spectrum of transcriptional states that are often distinct from those of macrophages in non-cancerous tissue [Wu et al., 2025, Das et al., 2024, Chi et al., 2024, Wen et al., 2024, Pirrello et al., 2024, Xu et al., 2022, Mantovani et al., 2017, Cassetta et al.]. Although the relative abundance of individual TAM states varies between cancers, the core transcriptional modules seem to be conserved across tumor types, providing a starting point for biomarker discovery and predicting patient responses to immunotherapy [Wei et al., 2024]. A pan-cancer atlas that profiled ≈400, 000 immune cells across 17 cancer types found that the balance of concurrent TAM states, rather than total macrophage number, predicts immunotherapy response [Coulton et al., 2024]. Resolving TAM composition at the individual cell level is therefore critical to understanding the response and resistance to immune checkpoint therapies.

This motivates predicting patient-level immunotherapy response directly from single-cell profiles of macrophage cells. Single-cell RNA foundation models offer a natural starting point, although several obstacles remain. Indeed, current approaches to patient-level prediction from single-cell data collapse each patient into a pseudobulk profile before classification, discarding the cell-state heterogeneity that atlas studies identify as predictive signal [Coulton et al., 2024]. Single-cell cohorts are also small and heavily imbalanced, a regime in which flexible models may overfit to the majority class. Finally, foundation model embeddings are not natively interpretable: individual embedding dimensions carry no gene-level meaning, so a model that predicts well over them does not by itself yield a biological hypothesis.

Multiple Instance Learning (MIL) is a weakly-supervised learning paradigm in which training labels are available only at the bag level, while predictions are made by aggregating multiple unlabeled instances within each bag [Dietterich et al., Ilse et al.]. It has proven particularly successful in histopathology image analysis, where Whole Slide Images (WSIs) can be represented as bags of image patches: a slide is labeled positive if it contains at least one cancerous region, but the precise location of malignant tissue is unknown during training [Campanella et al., Lu et al.]. Attention-based MIL architectures learn to automatically identify and weight the most diagnostically relevant patches, enabling interpretable predictions without expensive pixel-level annotations [Ilse et al., Shao et al.]. Single-cell transcriptomics data exhibits a strikingly analogous structure, and we show it can be treated the same way: each patient represents a bag containing thousands of individual cells (instances), with clinical outcome known only at the patient level. Just as WSI-MIL methods identify informative tissue regions, single-cell MIL can identify disease-relevant cell populations and states without celllevel labels, while attention weights indicate which cells drove each prediction.

Our work addresses the limitations above through three key contributions: (1) *Patient-level response prediction from foundation model embeddings*. We apply attention-based MIL to macrophage embeddings from a single-cell RNA foundation model and benchmark it against pseudobulk aggregation, PCA-mean, and embedding-mean baselines across three public anti-PD1 cohorts, showing that preserving cell-level granularity improves patient-level discrimination over aggregation-based alternatives. (2) *Weighted MIL pooling with a supervised contrastive objective*, which applies class-specific weights to attention scores and combines focal loss with supervised contrastive learning [Khosla et al.] to focus on minority class patterns; (3) *Gene-level interpretation of foundation model embeddings*: a post-hoc attribution pipeline that combines attention weights, integrated gradients, and gene–embedding correlation to rank genes by their contribution to the patient-level prediction, from which we recover the regulatory programs distinguishing responders from non-responders.

## 2 Methods

### 2.1 Single-cell Data Preprocessing

We used three publicly available single-cell RNA-seq datasets of pre- and post-anti-PD1 treatment cancer biopsies with treatment response metadata: a lung cancer cohort (Liu) [Liu et al.], and two breast cancer datasets - a reannotated version of Bassez et al. (Bassez) by Gondal et al. [a] and Shiao [Shiao et al.]. Accession numbers are available in section 5. For Shiao, response to therapy was defined using the Residual Cancer Burden (RCB) classes provided with the dataset, which range from 0 to 3, where 0 denotes pathologic complete response (best outcome) and 3 indicates extensive residual invasive disease (worst outcome). RCB classes 0 and 1 were grouped as *responders* (coded 1) and RCB classes 2 and 3 as *non-responders* (coded 0), reflecting clinically meaningful distinctions between minimal and substantial residual disease following therapy. Gene names were harmonized across datasets (section 2.1.1) and immune cells were identified and extracted (see section 2.1.3).

#### 2.1.1 Gene Name Harmonization

HGNC gene symbols were mapped to Ensembl v110 [Harrison et al.] by checking each symbol against five Cell Ranger versions [Zheng et al.] (GRCh38-2020-A, grch38_3_0_0, hg19_3_0_0, grch38_1_2_0 and hg19_1_2_0) and assigning it an Ensembl identity. Symbols were excluded if they were absent from these Cell Ranger versions, mapped to deprecated Ensembl identifiers, or mapped inconsistently across Cell Ranger versions (to prevent them from incorrectly being assigned the wrong identity). Counts were summed where a symbol mapped to multiple Ensembl IDs in release 110, and where multiple HGNC symbols converged to the same Ensembl ID. For Bassez, 8,291 genes were removed and 5,144 duplicates merged, leaving a total of 29,873 genes, for Shiao, 1,345 genes were removed and 31 duplicates merged and for Liu, 1,686 genes were removed and 94 duplicates merged, leaving 30,050 genes.

#### 2.1.2 MALAT1 Expression Filtering

An additional sample-wise MALAT1 quality control step was applied following the publicly available MALAT1 thresholding approach of Clarke and Bader (https://github.com/BaderLab/MALAT1_threshold). MALAT1 is highly expressed in the nucleus, and very low expression is characteristic of cytosolic debris and empty droplets [Clarke and Bader, Montserrat-Ayuso and Esteve-Codina]. MALAT1 counts were normalized to a library size of 10^4^, and an adaptive per-sample threshold was calculated from the density of normalized expression, following the Clarke and Bader algorithm (peak identification, local minima detection, and quadratic fitting around the principal MALAT1 peak). A minimum normalized MALAT1 expression threshold of 0.3 was used where the automated procedure failed to identify a reliable peak. Cells with MALAT1 expression at or below the inferred threshold were removed; high-MALAT1 cells were retained, as high MALAT1 expression can reflect metabolic activity rather than damage.

#### 2.1.3 Single-cell Dimensionality Reduction and Clustering

Single-cell expression counts were log-normalized to a library size of 10^4^, and highly variable genes were selected per dataset for principal component analysis. The technical variation between samples was corrected before downstream analysis when batch effects were evident. Nearest neighbor graphs were constructed in the resulting latent space, graph-based clustering was used to identify transcriptionally related cell populations and UMAPs were generated for visualization.

Immune cells were identified by combining the original cell-type annotations with immune cell expression profiles, assigned at both the cell and cluster levels, and verified against canonical marker genes.

Following MALAT1 filtering and immune cell selection, 102,174 cells remained for Bassez, 1,247,333 for Liu, and 3,405,015 for Shiao. Macrophages were then subclustered at higher resolution within each dataset, and macrophage subtypes assigned from differentially expressed marker genes. Cell embeddings were generated with the Biomedical Foundation Model (BMFM-RNA) [Danziger et al., 2026], which was also used to identify the myeloid compartment and further annotate macrophages.

### 2.2 Multiple Instance Learning with Supervised Contrastive Focal Loss

Inspired by the approach of Whole Slide Imaging in histology, we employed a Multiple Instance Learning (MIL) framework to aggregate single-cell macrophage representations into patient-level predictions. Each patient was treated as a “bag” containing multiple cell “instances”, and cell-level embeddings were generated using BMFM-RNA. The MIL architecture consisted of an attention-based pooling mechanism that learned to weight cells according to their importance for patient-level classification. Specifically, for a patient with *N* cells, let **x**_*i*_ ∈ ℝ^*d*^ denote the BMFM-RNA embedding of cell *i*, where *d* = 368 is the embedding dimension. The attention network computed importance scores *a*_*i*_ for each cell:

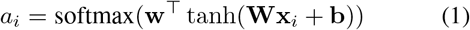

where **W** ∈ ℝ^*h×d*^ and **w** ∈ ℝ^*h*^ are learnable parameters with hidden dimension *h* = 128. The patient-level representation was obtained via weighted aggregation: 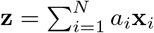.

To address class imbalance and improve discriminative feature learning, we incorporated focal loss [Lin et al.] with supervised contrastive learning [Khosla et al.]. Focal loss served as the primary classification objective, with focusing parameter *γ* = 2.0 and class-weighting parameter *α* = 0.25, to down-weight easy examples and concentrate learning on hard-to-classify patients - particularly critical in imbalanced clinical datasets. The focal loss is defined as:

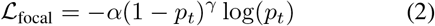

where *p*_*t*_ is the predicted probability for the true class and *α* is the class weight. The supervised contrastive loss encouraged patient representations from the same class to cluster together while pushing apart different classes. For a batch of patients, the contrastive loss for patient *i* is:

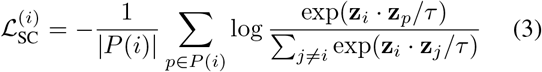

where *P* (*i*) denotes the set of positive pairs (same class as patient *i*), *j i* represents all other patients in the batch, and *τ* = 0.07 is the temperature parameter. The total loss combined both objectives:

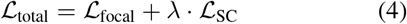

where *λ* = 0.5 balances the classification and contrastive objectives, and 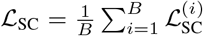 is averaged over the batch of size *B*. During training, focal loss was applied to logits from a linear classifier head, while contrastive loss operated on the normalized patient-level representations **z** before classification.

### 2.3 Benchmarking Framework

#### 2.3.1 Baseline Methods

We compared our approach against multiple patient-level aggregation strategies and MIL architectures.

##### Simple Aggregation Methods

These methods aggregate cell-level representations into fixed patient-level features: (1) *Pseudobulk + Random Forest*, which sums gene expression across all cells per patient and applies a Random Forest classifier; (2) *Mean PCA + Logistic Regression*, which averages PCA-reduced expression profiles across cells; and (3) *Mean/Median Embeddings + Logistic Regression*, which aggregates foundation model embeddings using mean or median operations followed by logistic regression.

##### Multiple Instance Learning Methods

MIL methods treat each patient as a bag of cell instances without requiring cell-level labels. We evaluated: (1) *Standard MIL variants* including attention-based pooling [Ilse et al.], gated attention [Ilse et al.], and max-pooling aggregation; (2) *Improved MIL* architectures with enhanced regularization optimized for small sample sizes; (3) *Weighted MIL*, which addresses class imbalance through instance-level reweighting; and (4) *Supervised Contrastive MIL (SC-MIL)* variants that combine attention mechanisms with supervised contrastive learning objectives [Khosla et al.] to improve representation learning and class separation. SC-MIL methods were evaluated with contrastive loss weights of 0.5 and 0.7 to assess robustness to class imbalance.

### 2.4 Model Training and Evaluation

All models were evaluated using repeated stratified 5-fold cross-validation with 10 repeats (50 total folds) to ensure robust performance estimates while maintaining class balance across folds. The MIL model was optimized using the Adam optimizer with learning rate 10^*™*3^ and weight decay 10^*™*4^, training for a maximum of 50 epochs with early stopping (patience=10 epochs). The combined loss function was minimized using mini-batch gradient descent with batch size of 1, as each patient constitutes a single bag in the MIL framework. After MIL training, patient-level representations were extracted and a logistic regression classifier was trained on these representations. Model performance was assessed using area under the ROC curve (AUROC), accuracy, weighted F1-score, and Matthews correlation coefficient (MCC), with statistical significance evaluated using Wilcoxon signed-rank tests and false discovery rate (FDR) correction for multiple comparisons.

### 2.5 Attention-Based Cell Importance Analysis

To identify which macrophage cells were most informative for treatment response prediction, we performed attention analysis on the trained MIL models. For each patient bag containing *N* cells, the attention mechanism assigned importance weights 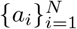 to individual cells, where 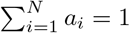. These weights quantified each cell’s contribution to the patient-level prediction, with higher weights indicating greater importance for classification. We write **a** = (*a*_1_, …, *a*_*N*_)^⊤^ ∈ ℝ^*N*^ for the vector of attention weights of a given patient bag.

We extracted the top *k* = 20 cells with highest attention weights per patient, representing the most discriminative cells for response prediction.

### 2.6 Gene Importance Extraction via Attention-Weighted Correlation

As it is not always possible or computationally efficient to retrieve feature-level information from Foundation Model (FM) embeddings, we implemented a post-hoc interpretation method combining attention, integrated gradients and correlation analysis to map embedding-level importance back to individual genes.

To understand *why* a given cell was important, we employed Integrated Gradients [Sundararajan et al.], which attributes a prediction to input features by integrating gradients along a path from a baseline (zero vector) **x**_b_ to the cell embedding **x**_i_:

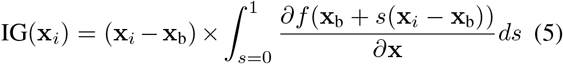

where *f* (*·*) represents the complete MIL model (aggregation and classifier), **x**_*i*_ is the BMFM-RNA embedding of cell *i*, **x**_b_ is the baseline, and the integral was approximated using the Riemann trapezoidal rule with *n* = 50 steps. This yields attribution scores IG(**x**_*i*_) ∈ ℝ^*d*^ indicating which embedding dimensions drove the model’s prediction for that cell.

For a given patient, let **X** ∈ ℝ^*N ×G*^ denote the gene expression matrix over *N* cells and *G* genes, and **E** ∈ ℝ^*N ×d*^ represent the corresponding cell embeddings. Combining the attention weights **a** of section 2.5 with integrated gradients gives the attention-weighted importance:

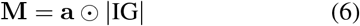

where ⊙ denotes element-wise multiplication and aggregating across cells yields per-embedding-dimension importance:

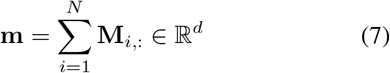

To map embedding-level importance to individual genes, we computed Pearson correlations between gene expression and embedding dimensions. For gene *k* and embedding dimension *j*, the correlation coefficient is:

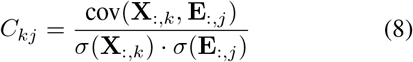

where cov(*·,·*) denotes covariance and *σ*(*·*) denotes standard deviation.

Gene importance scores were computed as a weighted sum of absolute correlations:

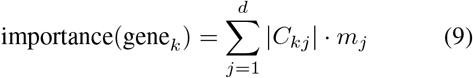

For each patient, genes were ranked by importance, and the top 100 genes were retained for downstream analysis.

### 2.7 Gene Regulatory Network Construction and Analysis

Because correlation alone is insufficient to interpret the resulting cell states, we performed gene regulatory network (GRN) analysis on the top-attended macrophage cells identified in section 2.5. These cells were aggregated by response class (responders vs. non-responders), and mean gene expression profiles were computed for each group.

Transcription factor (TF) activities were inferred using the Univariate Linear Model (ULM) method implemented in decoupler [Badia-i Mompel et al., 2022], with the CollecTRI prior-knowledge network [Müller-Dott et al., 2023] providing curated TF-target gene interactions. For a given response class *c*, let **E**_*c*_ *∈* ℝ^*G*^ denote the mean log-normalized expression of *G* genes, and let **w**_*t*_ = **R**_:,*t*_ *∈* ℝ^*G*^ denote the regulon of TF *t*, where *R*_*gt*_ represents the mode of regulation (activation: +1, repression: *™* 1, 0 if gene *g* is not a target of TF *t*). ULM treats the genes of a single observation as the sample population and fits an independent linear model for each TF, with the regulon weights as the sole covariate:

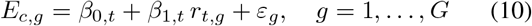

The activity of TF *t* is taken as the *t*-value of the fitted slope,

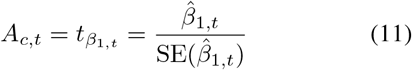

Positive values indicate coordinated up-regulation of the TF’s targets relative to the rest of the transcriptome, weighted by mode of regulation.

For visualization, we selected the top *n*_*s*_ = 10 TFs with highest absolute activity scores and their top *n*_*t*_ = 25 target genes with highest absolute expression. Networks were visualized using the Kamada-Kawai force-directed layout algorithm, with node colors representing TF activities (red-blue diverging colormap) and target gene expression levels (viridis colormap).

#### 2.7.1 Differential Network Analysis

To identify regulatory differences between response classes, we computed differential networks by subtracting non-responder from responder values:

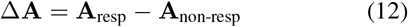

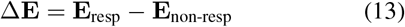

where Δ**A** represents differences in TF activities and Δ**E** represents differences in target gene expression. Positive values indicate higher activity or expression in responders. The differential network was visualized using a red-blue diverging colormap centered at zero, with red nodes and edges indicating up-regulation in responders and blue indicating up-regulation in non-responders. Hub TFs with large activity differences and many differentially expressed targets represent candidate regulatory modules associated with treatment response.

## 3 Results

### 3.1 Patient-level prediction of treatment response

We first asked whether macrophage embeddings carry sufficient signal to predict anti-PD1 response at the patient level, and whether preserving single-cell granularity offers any advantage over collapsing each patient into a single feature vector before classification. Ten strategies were evaluated under repeated stratified cross-validation: four aggregation baselines (pseudobulk with a random forest, PCA-mean, and mean or median embeddings with logistic regression) and six MIL configurations spanning standard attention, gated and improved gated attention, weighted pooling, and supervised contrastive variants (fig. 2).

**Figure 1.**
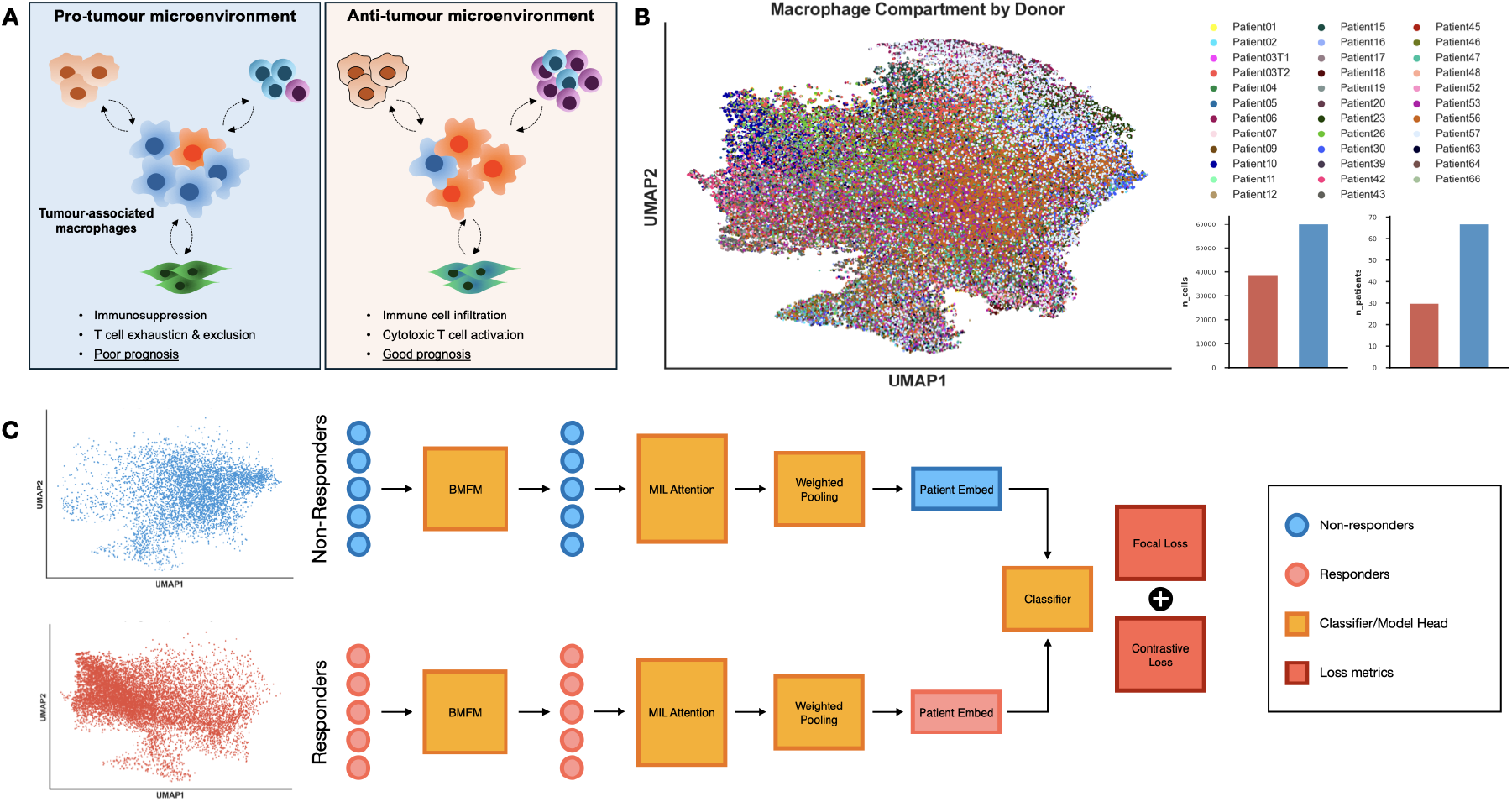
Experimental design and model architecture schematic. **A.** Schematic which describes an overview of the pro- and anti-tumor microenvironment driven by macrophages. **B**. UMAP of the BMFM-RNA embedding of the macrophage compartment of Shiao et al. [2024] with the number of cells and patients per condition. **C**. Model architecture and approach taken to train a MIL-attention network with supervised contrastive learning and focal loss for minimal data learning.

**Figure 2.**
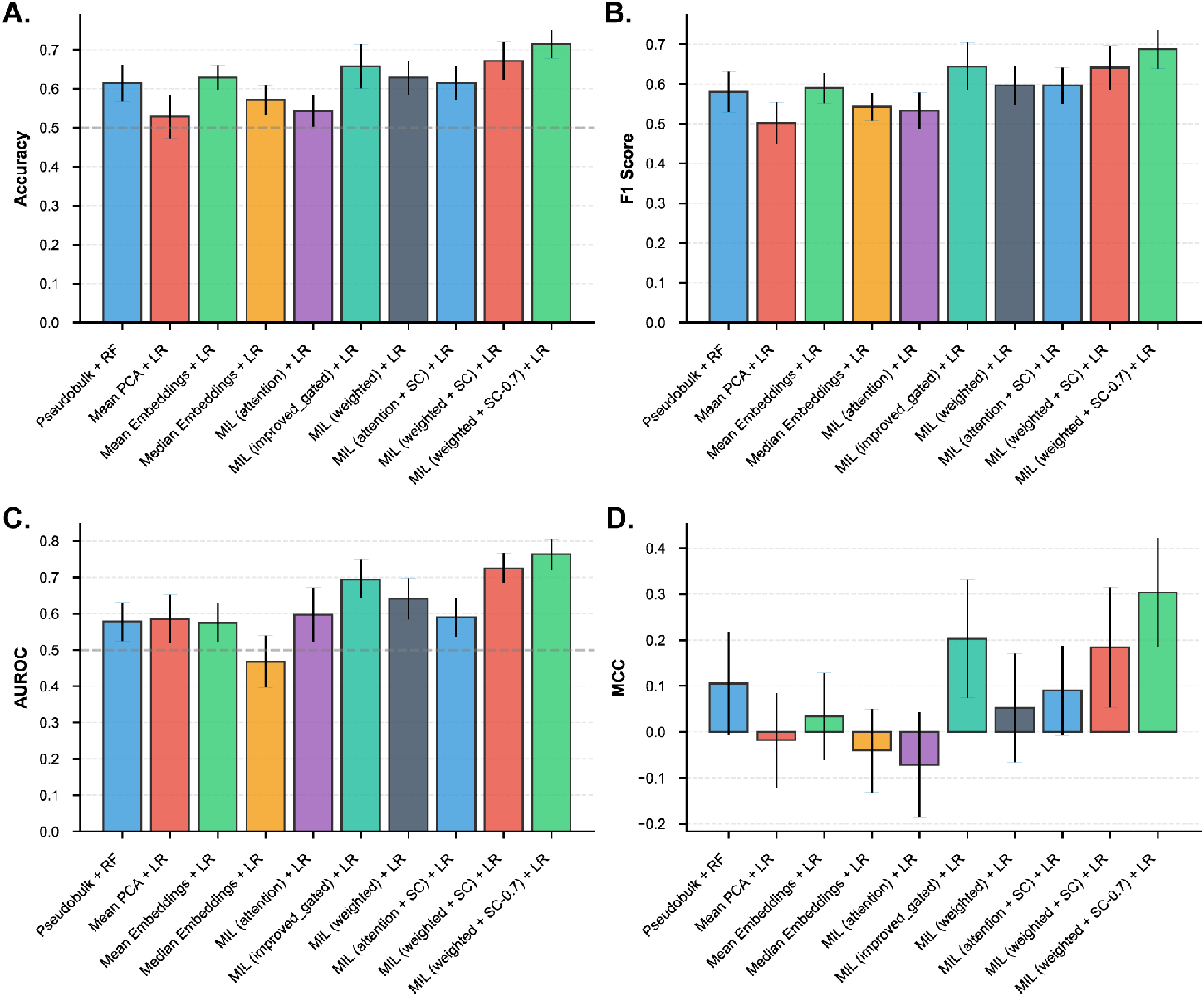
Predicting patient responders and non-responders at the patient-level. **(A,B,C)** Bar plots with standard deviation of the Accuracy, F1-score and Area Under the Receiver Operating Characteristic Curve (AUROC). **(D)** Bar plot with standard deviation of the Matthew’s Correlation Coefficient (MCC) where the higher (more positive) MCC values describe better agreement between predictions and the true labels, whereas negative values indicate performance below random expectation.

Methods retaining cell-level structure performed best. Pseudobulk with a random forest reached an AUROC of 0.58 and mean-embedding logistic regression 0.57, while the strongest MIL configuration, weighted pooling with a supervised contrastive objective at *λ* = 0.7, reached 0.77 (fig. 2C); the same ordering held for accuracy, weighted F1 and MCC (fig. 2A,B,D). Differences across methods were significant by Friedman test for accuracy (*p* = 0.048) and AUROC (*p* = 0.027), but not for weighted F1 (*p* = 0.072) or MCC (*p* = 0.092), so we treat the former two as the primary evidence.

The improvement appears attributable to the dual-objective formulation rather than to MIL alone: standard attention MIL performed comparably to the aggregation baselines, with gains emerging specifically in the weighted-pooling and supervised contrastive variants (fig. 2C). This is consistent with the combined loss allowing the model to optimize simultaneously for discriminative class boundaries (via focal loss) and for a semantically meaningful representation geometry (via contrastive learning). Absolute performance nonetheless remains modest, with MCC intervals extending toward zero (fig. 2D) as expected at this sample size. We therefore read these results as evidence that SC-MIL recovers a macrophage-restricted signal predictive of patient-level immunotherapy response.

### 3.2 Contribution of focal and contrastive loss terms

To systematically evaluate how each loss component contributes to MIL performance, we conducted an ablation comparing three loss configurations: focal loss only, supervised contrastive loss only, and their combination. Each configuration was tested across three MIL architectures (with focal loss only, contrastive loss only and combined focal and contrastive) on synthetic datasets with varying class imbalance ratios (50%, 30%, 20%, 10%, and 5% minority class). Synthetic data was used here because it allows the minority-class fraction to be varied systematically, which the clinical cohorts do not permit.

The combined objective outperformed either loss function alone across all MIL architectures and imbalance levels (Figure 3). For the attention-based MIL architecture at 20% minority class, it achieved a mean accuracy of 0.847*±*0.023 (mean *±* SD), compared to 0.782*±*0.031 for focal loss only and 0.756 *±* 0.028 for contrastive loss only, with the same ordering for weighted F1 (0.823 *±* 0.027; 0.751 *±* 0.034; 0.728 *±* 0.031) and MCC (0.694 *±* 0.046; 0.564 *±* 0.052; 0.512 *±* 0.048).

**Figure 3.**
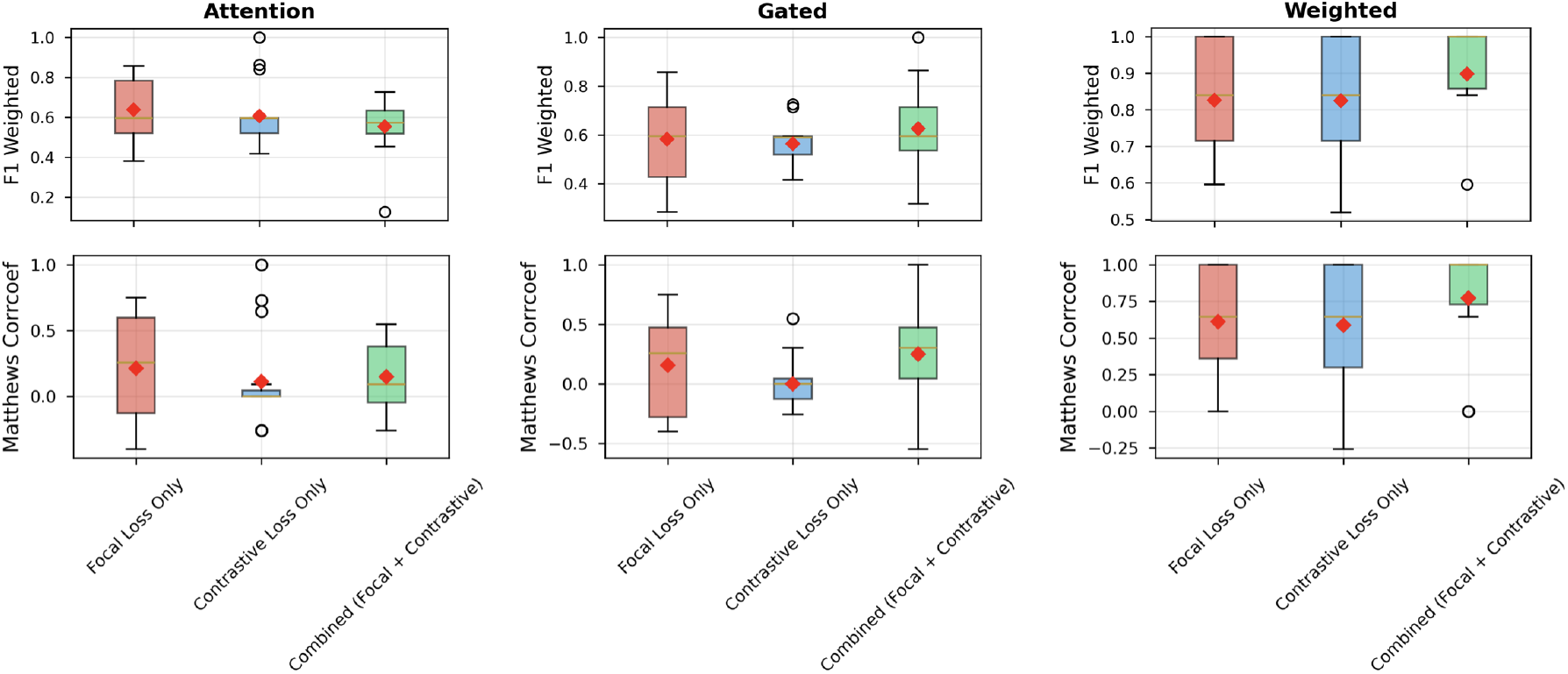
Ablation study investigating different MIL attention model with differing loss metrics for imbalanced datasets.

The advantage widens with imbalance. At 5% minority class the combined loss reached an AUROC of 0.912 ± 0.018 against 0.834 *±* 0.027 for focal loss only (*p <* 0.001, *d* = 3.45) and 0.798 *±* 0.031 for contrastive loss only (*p <* 0.001, *d* = 4.23). Accuracy for contrastive loss alone degraded significantly (0.623 *±* 0.041), but remained stable for the combined loss (0.831 *±* 0.024). This is consistent with the two terms addressing different failure modes: focal loss rebalances the classification gradient towards hard-to-classify minority patients, while contrastive learning shapes the geometry of the patient representation. Across all 75 conditions, the combined approach ranked first in 68 cases for accuracy and 71 for MCC, indicating the benefit is not specific to a particular architecture.

### 3.3 Gene-level attribution recovers a coherent myeloid program

The trained model assigns each cell in a patient bag a weight reflecting its contribution to the patient-level prediction (section 2.5), so we first used these weights to select the 20 most-attended macrophages per patient, giving a set of cells the model itself identified as most discriminative for response. We then applied the post-hoc attribution analysis described in section 2.6 to these cells, combining attention with integrated gradients and gene-embedding correlation to interpret the predictions at gene resolution.

The resulting ranking (fig. 4) is dominated by a coherent mononuclear-phagocyte program. The top-ranked gene, FCN1, marks monocyte-derived cells recently recruited to the tumor microenvironment: FCN1^+^ TAMs act as transitional monocyte-like precursors capable of differentiating into more specialized subsets [Wang et al., 2024], and have been implicated in chemotherapy response in other cancer types [Xi et al., 2023]. Several highly ranked genes have established TAM associations: CD63 and APOC1 are highly expressed in TAMs, where they promote proliferation, angiogenesis and poor prognosis across cancers [**?**]. Complement and lysosomal components follow (C1QA, CTSB, CTSD, PSAP, CSTB), alongside the pan-macrophage marker CD68, the lectin CLEC10A, and GPNMB, which marks a lipid-associated TAM state [Dang Cao et al., 2025].

**Figure 4.**
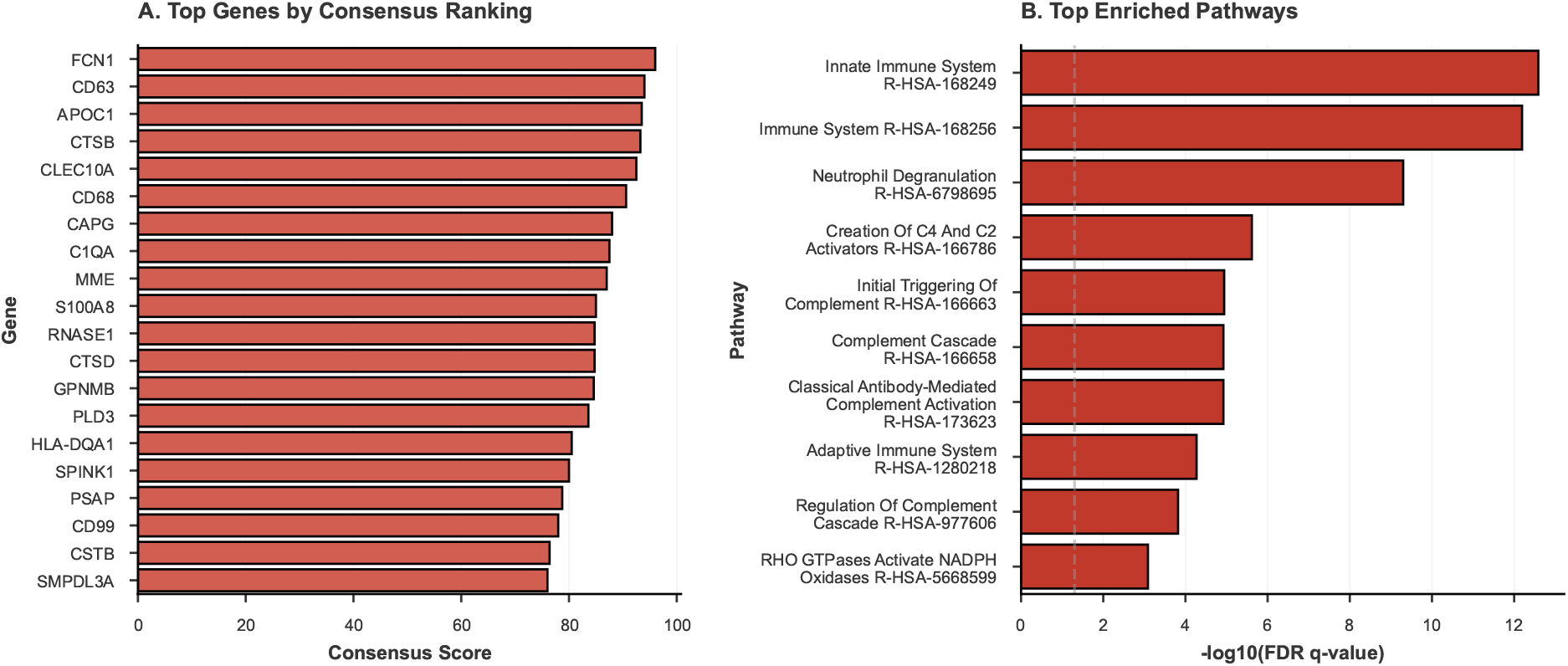
Post-hoc gene importance scores using attention-weighted correlation. **(A)** Top responder-associated marker genes, ranked by consensus scores across patients. **(B)** Pathways most enriched among top genes. The top 50 ranked genes were used as input for over-representation analysis based on the Reactome Pathway database.

Over-representation analysis of the top 50 ranked genes against Reactome (fig. 4B) returned innate immune signaling, neutrophil degranulation and a block of complement cascade terms. Because the ranking derives from correlations between genes and embedding dimensions, it is likely to be biased towards genes with high expression and high variance across cells; indeed, several ranked genes (CTSB, PSAP, CSTB) are abundant in most myeloid cells.

### 3.4 Regulatory program separate responders from non-responders

To understand the mechanism underlying the model’s predictions treatment response, we inferred transcription-factor activity on the top-attended cells from each patient, aggregated by response class. The two classes yielded structurally distinct networks (fig. 5). Macrophages from non-responder samples showed increased activity of E2F1 and ETV4, the latter a known activator of hypoxia-inducible factor signaling [Wollenick et al.]. In contrast, macrophages from responder samples showed higher activity across two coherent modules: antigen presentation (CIITA, RFX5, RFXAP, RFXANK) and the response to interferon (STAT1, IRF1, IRF9). The differential network separates them cleanly (fig. 5C).

**Figure 5.**
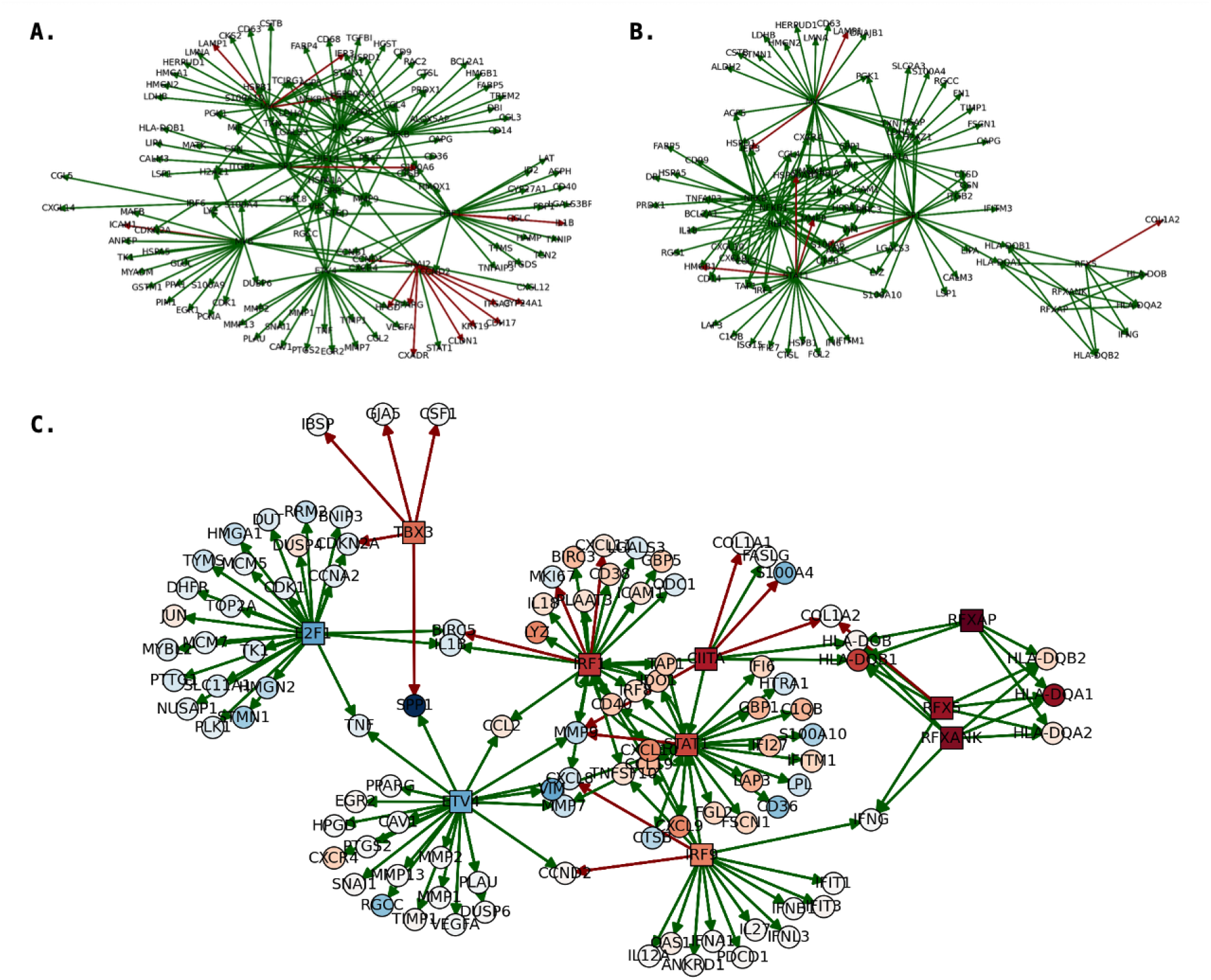
Consensus macrophage signature between responders and non-responders. **(A)** TF-Gene regulatory networks representing the aggregated signature of the top cells from each patient in the non-responders, while **(B)** represents the top cells from the responders. **(C)** Differential network analysis of the responders and non-responders. Blue nodes represent up-regulation in the non-responders and red nodes show up-regulation in the responders.

This hypoxia-versus-interferon contrast maps onto an established axis of TAM polarity. The CXCL9:SPP1 ratio defines a polarity axis based on the balance between CXCL9 (pro-inflammatory, anti-tumour) and SPP1 (osteopontin, pro-tumour) expression, and is an independent prognostic marker across cancer types [Bill et al., a]. Notably, it is independent of classical M1/M2 markers, which often fail to predict clinical outcomes or immune context *in vivo* [Qian and Pollard].

Our model recovers this axis without being supervised towards it. SPP1 is up-regulated in the non-responder network, while CXCL9 is up-regulated within the responder interferon module (fig. 5C), reconstructing, reconstructing the CS axis from patient-level response labels alone. Neither gene was supplied as a feature, label or filter at any stage, which is strong evidence that the attention mechanism selects biologically meaningful cells rather than fitting cohort-specific noise. Moreover, because the networks are built from top-attended cells only, they identify the regulatory context in which CXCL9 and SPP1 sit: the transcription factors and neighboring targets co-active with each axis member in the cells the model found most predictive. This context is not recoverable from bulk or pseudobulk measurement of the CXCL9:SPP1 ratio itself.

## 4 Discussion

In this study, we leverage approaches from NLP and image-based machine learning to predict patient-level responses to anti-PD-1 treatment from single cell RNA-seq data. Anti-PD-1/PD-L1 (programmed death 1 or programmed death ligand 1) checkpoint inhibitor therapies work well in only ~ 30% of patients [Ribas and Wolchok]. Effective anti-PD(L)1 responses require cytotoxic CD8^+^ T-cell recruitment into the tumor; however, these responses can be impaired by T-cell exhaustion, T-cell exclusion, and an immunosuppressive tumor microenvironment (TME) [Jiang et al.]. CD8^+^ T-cell infiltration is regulated by the CXCR3 chemokine pathway [House et al., Dangaj et al.], whose ligands CXCL9 and CXCL10 are produced by macrophages and dendritic cells, contributing to a “hot” TME permissive to checkpoint blockade [Reschke and Gajewski, Marcovecchio et al.]. Response is therefore not determined by any single cell type or marker but by the coordinated state of the microenvironment, which makes it difficult to predict from bulk measurements yet well suited to single-cell approaches that can resolve the contributing states individually.

Using our MIL-based framework, we recover established regulators of anti-PD-1 response at the gene level and identify differences in the regulatory networks of TAMs between responders and non-responders, taking advantage of the granularity provided by single-cell data by using cell-level embeddings from a single-cell foundation model rather than pseudobulk aggregates as model input.

In particular, we recovered differences along the CXCL9:SPP1 (CS) ratio, an independent prognostic marker across cancer types. A high ratio correlates with improved overall survival and better response to anti-PD-1 immunotherapy, while a low ratio correlates with poor prognosis [Bill et al., b]. The two markers label largely mutually exclusive phenotypes: the SPP1-linked phenotype is induced in hypoxic environments, while CXCL9 is associated with IFN-high environments, and IFN*γ* is a master regulator of CXCL9 expression [Tokunaga et al.]. This is consistent with the upregulation of CXCR3 ligands following checkpoint blockade [House et al.]. Notably, the axis is independent of classical M1/M2 markers, which often fail to predict clinical outcomes or immune context *in vivo*, and TAMs comprise multiple transcriptional states capable of expressing either gene to varying degrees, indicating plasticity rather than fixed polarization. CXCL9 alone is an unreliable pan-cancer prognostic marker, being associated with better outcomes in some settings and worse in others [Ding et al.]; it is the balance between CXCL9^+^ and SPP1^+^ subpopulations that carries prognostic weight.

The advantage of the CS ratio is that it is simple and measurable by histology or qPCR, capturing TME-wide coordination in a single quantity. However, that signal can be diluted in noisy single-cell data, and the ratio alone says nothing about the regulatory context in which either gene sits. Because our networks are constructed from the cells the model itself found most predictive, they identify the transcription factors and first-neighbor targets co-active with each axis member, suggesting genes that might be targeted alongside it. More broadly, this MIL framework enables cell-specific analyses rather than relying on a bulk approach, providing cellular-level insight into the role of the CXCL9/SPP1 axis in TAMs.

More generally, our results suggest that incorporating the full spectrum of macrophage subpopulations and their transcriptional states into prognostic models, rather than single markers or ratios, could improve both predictive accuracy and biological insight, particularly in cancers where CXCL9 expression alone is neutral or associated with worse outcomes.

## Limitations

There are some trade-offs and limitations to this approach. Firstly, it relies on linear correlation, which may not capture complex non-linear relationships between genes and embeddings, and genes may correlate with important embeddings for reasons unrelated to the prediction task. Additionally, as it is also a post-hoc method, it infers gene importance indirectly rather than directly from the model’s decision process. Finally, computing correlations on a per-patient basis with limited cells may lead to unstable estimates. Despite these limitations, this method provides interpretable gene-level insights from models operating on learned embeddings, enabling biological validation and hypothesis generation.

## 5 Data and Code Availability

The publicly available datasets used in this study are accessible in GEO accessions: Liu lung cancer (GEO: GSE243013 [Liu et al.]); Shiao breast cancer (GEO: GSE246613 [Shiao et al.]); a reannotated version of Bassez et al. breast cancer dataset (merged_tissue_cluster_update_filt_other_mal_4.RDS object from Zenodo Gondal et al. [b]).

## References

Emmanuel L. Gautier, Tal Shay, Jennifer Miller, Melanie Greter, Claudia Jakubzick, Stoyan Ivanov, Julie Helft, Andrew Chow, Kutlu G. Elpek, Simon Gordonov, Amin R. Mazloom, Avi Ma’ayan, Wei-Jen Chua, Ted H. Hansen, Shannon J. Turley, Miriam Merad, and Gwendalyn J. Randolph. Gene-expression profiles and transcriptional regulatory pathways that underlie the identity and diversity of mouse tissue macrophages. Nature Immunology, 13(11):1118–1128, November 2012. ISSN 1529-2916. doi:10.1038/ni.2419. URL https://www.nature.com/articles/ni.2419. Publisher: Nature Publishing Group.

D A Hume. Differentiation and heterogeneity in the mononuclear phagocyte system. Mucosal Immunology, 1(6):432–441, November 2008. ISSN 1933-0219. doi:10.1038/mi.2008.36. URL https://www.sciencedirect.com/science/article/pii/S1933021922017032.

Siamon Gordon and Philip R. Taylor. Monocyte and macrophage heterogeneity. Nature Reviews Immunology, 5(12):953–964, December 2005. ISSN 1474-1741. doi:10.1038/nri1733. URL https://www.nature.com/articles/nri1733. Publisher: Nature Publishing Group.

Marie Laviron and Alexandre Boissonnas. Ontogeny of Tumor-Associated Macrophages. 10. ISSN 1664-3224. doi:10.3389/fimmu.2019.01799. URL https://www.frontiersin.org/journals/immunology/articles/10.3389/fimmu.2019.01799/full.

Qiong-wen Zhang, Lei Liu, Chang-yang Gong, Hua-shan Shi, Yun-hui Zeng, Xiao-ze Wang, Yu-wei Zhao, and Yu-quan Wei. Prognostic Significance of Tumor-Associated Macrophages in Solid Tumor: A Meta-Analysis of the Literature. 7(12):e50946. ISSN 1932-6203. doi:10.1371/journal.pone.0050946. URL https://journals.plos.org/plosone/article?id=10.1371/journal.pone.0050946.

Bin-Zhi Qian and Jeffrey W. Pollard. Macrophage Diversity Enhances Tumor Progression and Metastasis. Cell, 141(1):39–51, April 2010. ISSN 0092-8674, 1097-4172. doi:10.1016/j.cell.2010.03.014. URL https://www.cell.com/cell/abstract/S0092-8674(10)00287-4. Publisher: Elsevier.

Thomas A. Wynn, Ajay Chawla, and Jeffrey W. Pollard. Macrophage biology in development, homeostasis and disease. Nature, 496(7446):445–455, April 2013. ISSN 1476-4687. doi:10.1038/nature12034. URL https://www.nature.com/articles/nature12034. Publisher: Nature Publishing Group.

Jacob Stultz and Lawrence Fong. How to turn up the heat on the cold immune microenvironment of metastatic prostate cancer. Prostate Cancer and Prostatic Diseases, 24(3):697–717, September 2021. ISSN 1476-5608. doi:10.1038/s41391-021-00340-5. URL https://www.nature.com/articles/s41391-021-00340-5. Publisher: Nature Publishing Group.

Ann Erlandsson, Jessica Carlsson, Marie Lundholm, Anna Fält, Sven-Olof Andersson, Ove Andrén, and Sabina Davidsson. M2 macrophages and regulatory T cells in lethal prostate cancer. The Prostate, 79(4):363–369, 2019. ISSN 1097-0045. doi:10.1002/pros.23742. URL https://onlinelibrary.wiley.com/doi/abs/10.1002/pros.23742. _eprint: https://onlinelibrary.wiley.com/doi/pdf/10.1002/pros.23742Y.

Marie Lundholm, Christina Hägglöf, Maria L. Wikberg, Pär Stattin, Lars Egevad, Anders Bergh, Pernilla Wikström, Richard Palmqvist, and Sofia Edin. Secreted Factors from Colorectal and Prostate Cancer Cells Skew the Immune Response in Opposite Directions. Scientific Reports, 5:15651, October 2015. ISSN 2045-2322. doi:10.1038/srep15651.

M. Lanciotti, L. Masieri, M. R. Raspollini, A. Minervini, A. Mari, G. Comito, E. Giannoni, M. Carini, P. Chiarugi, and S. Serni.The Role of M1 and M2 Macrophages in Prostate Cancer in relation to Extracapsular Tumor Extension and Biochemical Recurrence after Radical Prostatectomy. BioMed Research International, 2014:486798, 2014. ISSN 2314-6133. doi:10.1155/2014/486798. URL https://www.ncbi.nlm.nih.gov/pmc/articles/PMC3967497/.

Frances R. Balkwill and Alberto Mantovani. Cancer-related inflammation: Common themes and therapeutic opportunities. Seminars in Cancer Biology, 22 (1):33–40, February 2012. ISSN 1044-579X. doi:10.1016/j.semcancer.2011.12.005. URL https://www.sciencedirect.com/science/article/pii/S1044579X11001064.

Alberto Mantovani, Paola Allavena, Federica Marchesi, and Cecilia Garlanda. Macrophages as tools and targets in cancer therapy. Nature Reviews Drug Discovery, 21(11):799–820, November 2022. ISSN 1474-1784. doi:10.1038/s41573-022-00520-5. URL https://www.nature.com/articles/s41573-022-00520-5. Publisher: Nature Publishing Group.

Jiasheng Xu, Lei Ding, Jianfeng Mei, Yeting Hu, Xiangxing Kong, Siqi Dai, Tongtong Bu, Qian Xiao, and Kefeng Ding. Dual roles and therapeutic targeting of tumor-associated macrophages in tumor microenvironments. 10(1):268. ISSN 2059-3635. doi:10.1038/s41392-025-02325-5. URL https://www.nature.com/articles/s41392-025-02325-5.

Hans Prenen and Massimiliano Mazzone. Tumor-associated macrophages: A short compendium. 76(8): 1447–1458. ISSN 1420-9071. doi:10.1007/s00018-018-2997-3. URL https://doi.org/10.1007/s00018-018-2997-3.

Qiyao Yang, Ningning Guo, Yi Zhou, Jiejian Chen, Qichun Wei, and Min Han. The role of tumor-associated macrophages (TAMs) in tumor progression and relevant advance in targeted therapy. 10(11):2156–2170. ISSN 2211-3835. doi:10.1016/j.apsb.2020.04.004. URL https://www.sciencedirect.com/science/article/pii/S2211383520305475.

Daniela F. Quail and Johanna A. Joyce. Microenvironmental regulation of tumor progression and metastasis. 19(11):1423–1437. ISSN 1546-170X. doi:10.1038/nm.3394. URL https://www.nature.com/articles/nm.3394.

ao Sun, Michael F. Cronin, Monique C. P. Mendonça, Jianfeng Guo, and Caitriona M. O’Driscoll. Sialic acid-targeted cyclodextrin-based nanoparticles deliver CSF-1R siRNA and reprogram tumour-associated macrophages for immunotherapy of prostate cancer. European Journal of Pharmaceutical Sciences, 185:106427, June 2023. ISSN 0928-0987. doi:10.1016/j.ejps.2023.106427. URL https://www.sciencedirect.com/science/article/pii/S0928098723000581.

Fangzhou Dou, Daoran Lu, and Jianjun Gao. Vimseltinib: A novel colony stimulating factor 1 receptor (CSF1R) inhibitor approved for treatment of tenosynovial giant cell tumors (TGCTs). 14(2):143–144. ISSN 2186-3644. doi:10.5582/irdr.2025.01010. URL https://pmc.ncbi.nlm.nih.gov/articles/PMC12143222/.

Vinod Nadella and Anu Sharma. Targeting Macrophages in Immunotherapy: The Ascent of CAR-Macrophages. 27(3):1292. ISSN 1422-0067. doi:10.3390/ijms27031292. URL https://pmc.ncbi.nlm.nih.gov/articles/PMC12897757/.

Renée Bouwstra, prefix=van useprefix=true family=Meerten, given=Tom, and Edwin Bremer. CD47-SIRP blocking-based immunotherapy: Current and prospective therapeutic strategies. 12(8):e943. ISSN 2001-1326. doi:10.1002/ctm2.943. URL https://pmc.ncbi.nlm.nih.gov/articles/PMC9339239/.

Lili Wu, Feihong Liang, Changgan Chen, Yaxin Zhang, Heguang Huang, and Yu Pan. Identification of prognostic and therapeutic biomarkers associated with macrophage and lipid metabolism in pancreatic cancer. Scientific Reports, 15(1):14584, April 2025. ISSN 2045-2322. doi:10.1038/s41598-025-99144-z. URL https://www.nature.com/articles/s41598-025-99144-z. Publisher: Nature Publishing Group.

Sreyashi Das, Mohan Kumar Dey, Ram Devireddy, and Manas Ranjan Gartia. Biomarkers in Cancer Detection, Diagnosis, and Prognosis. Sensors, 24(1):37, January 2024. ISSN 1424-8220. doi:10.3390/s24010037. URL https://www.mdpi.com/1424-8220/24/1/37. Number: 1 Publisher: Multidisciplinary Digital Publishing Institute.

Jianhua Chi, Qinglei Gao, and Dan Liu. Tissue-Resident Macrophages in Cancer: Friend or Foe? Cancer Medicine, 13(21):e70387, 2024. ISSN 2045-7634. doi:10.1002/cam4.70387. URL https://onlinelibrary.wiley.com/doi/abs/10.1002/cam4.70387. _eprint: https://onlinelibrary.wiley.com/doi/pdf/10.1002/cam4.70387.

Shaodi Wen, Renrui Zou, Xiaoyue Du, Rongtian Pan, Rutao Li, Jingwei Xia, Cong Xu, Ruotong Wang, Feng Jiang, Guoren Zhou, Jifeng Feng, Miaolin Zhu, Xin Wang, and Bo Shen. Identification of macrophage-related genes correlated with prognosis and immunotherapy efficacy in non-small cell lung cancer. Heliyon, 10(6):e27170, March 2024. ISSN 2405-8440. doi:10.1016/j.heliyon.2024.e27170. URL https://www.sciencedirect.com/science/article/pii/S2405844024032018.

Anthony Pirrello, Murray Killingsworth, Kevin Spring, John E. J. Rasko, and Dannel Yeo. Cancer-associated macrophage-like cells as a prognostic biomarker in solid tumors. The Journal of Liquid Biopsy, 6:100275, December 2024. ISSN 2950-1954. doi:10.1016/j.jlb.2024.100275. URL https://www.sciencedirect.com/science/article/pii/S2950195424001413.

Linping Xu, Meimei Yan, Jianpeng Long, Mengmeng liu, Hui Yang, and Wei Li. Identification of macrophage correlated biomarkers to predict the prognosis in patients with intrahepatic cholangiocarcinoma. Frontiers in Oncology, 12:967982, September 2022. ISSN 2234-943X. doi:10.3389/fonc.2022.967982. URL https://www.ncbi.nlm.nih.gov/pmc/articles/PMC9497456/.

Alberto Mantovani, Federica Marchesi, Alberto Malesci, Luigi Laghi, and Paola Allavena. Tumour-associated macrophages as treatment targets in oncology. Nature Reviews Clinical Oncology, 14(7):399–416, July 2017. ISSN 1759-4782. doi:10.1038/nrclinonc.2016.217. URL https://www.nature.com/articles/nrclinonc.2016.217. Publisher: Nature Publishing Group.

Luca Cassetta, Stamatina Fragkogianni, Andrew H. Sims, Agnieszka Swierczak, Lesley M. Forrester, Hui Zhang, Daniel Y. H. Soong, Tiziana Cotechini, Pavana Anur, Elaine Y. Lin, Antonella Fidanza, Martha Lopez-Yrigoyen, Michael R. Millar, Alexandra Urman, Zhichao Ai, Paul T. Spellman, E. Shelley Hwang, J. Michael Dixon, Lisa Wiechmann, Lisa M. Coussens, Harriet O. Smith, and Jeffrey W. Pollard. Human Tumor-Associated Macrophage and Monocyte Transcriptional Landscapes Reveal Cancer-Specific Reprogramming, Biomarkers, and Therapeutic Targets. 35(4):588–602.e10. ISSN 1535-6108, 1878-3686. doi:10.1016/j.ccell.2019.02.009. URL https://www.cell.com/cancer-cell/abstract/S1535-6108(19)30104-7.

Chen Wei, Yijie Ma, Mengyu Wang, Siyi Wang, Wenyue Yu, Shuailei Dong, Wenying Deng, Liangyu Bie, Chi Zhang, Wei Shen, Qingxin Xia, Suxia Luo, and Ning Li. Tumor-associated macrophage clusters linked to immunotherapy in a pan-cancer census. npj Precision Oncology, 8(1):1–20, August 2024. ISSN 2397-768X. doi:10.1038/s41698-024-00660-4. URL https://www.nature.com/articles/s41698-024-00660-4. Publisher: Nature Publishing Group.

Alexander Coulton, Jun Murai, Danwen Qian, Krupa Thakkar, Claire E. Lewis, and Kevin Litchfield. Using a pan-cancer atlas to investigate tumour associated macrophages as regulators of immunotherapy response. Nature Communications, 15(1):5665, July 2024. ISSN 2041-1723. doi:10.1038/s41467-024-49885-8. URL https://www.nature.com/articles/s41467-024-49885-8. Publisher: Nature Publishing Group.

Thomas G. Dietterich, Richard H. Lathrop, and Tomás Lozano-Pérez. Solving the multiple instance problem with axis-parallel rectangles. 89(1):31–71. ISSN 0004-3702. doi:10.1016/S0004-3702(96)00034-3. URL https://www.sciencedirect.com/science/article/pii/S0004370296000343.

Maximilian Ilse, Jakub Tomczak, and Max Welling. Attention-based deep multiple instance learning. In Proceedings of the 35th International Conference on Machine Learning, pages 2127–2136. PMLR. URL https://proceedings.mlr.press/v80/ilse18a.html.

Gabriele Campanella, Matthew G. Hanna, Luke Geneslaw, Allen Miraflor, Vitor Werneck Krauss Silva, Klaus J. Busam, Edi Brogi, Victor E. Reuter, David S. Klimstra, and Thomas J. Fuchs. Clinical-grade computational pathology using weakly supervised deep learning on whole slide images. 25(8):1301–1309. ISSN 1546-170X. doi:10.1038/s41591-019-0508-1. URL https://www.nature.com/articles/s41591-019-0508-1.

Ming Y. Lu, Drew F. K. Williamson, Tiffany Y. Chen, Richard J. Chen, Matteo Barbieri, and Faisal Mahmood. Data-efficient and weakly supervised computational pathology on whole-slide images. 5(6): 555–570. ISSN 2157-846X. doi:10.1038/s41551-020-00682-w. URL https://www.nature.com/articles/s41551-020-00682-w.

Zhuchen Shao, Hao Bian, Yang Chen, Yifeng Wang, Jian Zhang, Xiangyang Ji, and yongbing zhang. TransMIL: Transformer based correlated multiple instance learning for whole slide image classification. In Advances in Neural Information Processing Systems, volume 34, pages 2136–2147. Curran Associates, Inc. URL https://proceedings.neurips.cc/paper/2021/hash/10c272d06794d3e5785d5e7c5356e9ff-Abstract.html.

Prannay Khosla, Piotr Teterwak, Chen Wang, Aaron Sarna, Yonglong Tian, Phillip Isola, Aaron Maschinot, Ce Liu, and Dilip Krishnan. Supervised contrastive learning. In Advances in Neural Information Processing Systems, volume 33, pages 18661–18673. Curran Associates, Inc. URL https://proceedings.neurips.cc/paper/2020/hash/d89a66c7c80a29b1bdbab0f2a1a94af8-Abstract.html.

Stephen L. Shiao, Kenneth H. Gouin, Nathan Ing, Alice Ho, Reva Basho, Aagam Shah, Richard H. Mebane, David Zitser, Andrew Martinez, Natalie-Ya Mevises, Bassem Ben-Cheikh, Regina Henson, Monica Mita, Philomena McAndrew, Scott Karlan, Armando Giuliano, Alice Chung, Farin Amersi, Catherine Dang, Heather Richardson, Wonwoo Shon, Farnaz Dadmanesh, Michele Burnison, Amin Mirhadi, Zachary S. Zumsteg, Rachel Choi, Madison Davis, Joseph Lee, Dustin Rollins, Cynthia Martin, Negin H. Khameneh, Heather McArthur, and Simon R. V. Knott. Single-cell and spatial profiling identify three response trajectories to pembrolizumab and radiation therapy in triple negative breast cancer. Cancer Cell, 42(1):70–84.e8, January 2024. ISSN 1878-3686. doi:10.1016/j.ccell.2023.12.012.

Zedao Liu, Zhenlin Yang, Junqi Wu, Wenjie Zhang, Yuxuan Sun, Chao Zhang, Guangyu Bai, Li Yang, Hongtao Fan, Yawen Chen, Lei Zhang, Benyuan Jiang, Xiaoyan Liu, Xiaoshi Ma, Wei Tang, Chang Liu, Yang Qu, Lixu Yan, Deping Zhao, Yilong Wu, Shun He, Long Xu, Lishan Peng, Xiaowei Chen, Bolun Zhou, Liang Zhao, Zhangyi Zhao, Fengwei Tan, Wanting Zhang, Dingcheng Yi, Xiangjie Li, Qianqian Gao, Guangjian Zhang, Yongjie Wang, Minglei Yang, Honghao Fu, Yongjun Guo, Xueda Hu, Qingyuan Cai, Lu Qi, Yufei Bo, Hui Peng, Zhigang Tian, Yunlang She, Chang Zou, Linnan Zhu, Sijin Cheng, Yi Zhang, Wenzhao Zhong, Chang Chen, Shugeng Gao, and Zemin Zhang. A single-cell atlas reveals immune heterogeneity in anti-PD-1-treated non-small cell lung cancer. 188(11):3081–3096.e19. ISSN 0092-8674, 1097-4172. doi:10.1016/j.cell.2025.03.018. URL https://www.cell.com/cell/abstract/S0092-8674(25)00291-0.

Ayse Bassez, Hanne Vos, Laurien Van Dyck, Giuseppe Floris, Ingrid Arijs, Christine Desmedt, Bram Boeckx, Marlies Vanden Bempt, Ines Nevelsteen, Kathleen Lambein, Kevin Punie, Patrick Neven, Abhishek D. Garg, Hans Wildiers, Junbin Qian, Ann Smeets, and Diether Lambrechts. A single-cell map of intratumoral changes during anti-PD1 treatment of patients with breast cancer. 27(5):820–832. ISSN 1546-170X. doi:10.1038/s41591-021-01323-8. URL https://www.nature.com/articles/s41591-021-01323-8.

Mahnoor N. Gondal, Marcin Cieslik, and Arul M. Chinnaiyan. Integrated cancer cell-specific single-cell RNA-seq datasets of immune checkpoint blockade-treated patients. 12(1):139, a. ISSN 2052-4463. doi:10.1038/s41597-025-04381-6. URL https://www.nature.com/articles/s41597-025-04381-6.

Stephen L. Shiao, Kenneth H. Gouin, Nathan Ing, Alice Ho, Reva Basho, Aagam Shah, Richard H. Mebane, David Zitser, Andrew Martinez, Natalie-Ya Mevises, Bassem Ben-Cheikh, Regina Henson, Monica Mita, Philomena McAndrew, Scott Karlan, Armando Giuliano, Alice Chung, Farin Amersi, Catherine Dang, Heather Richardson, Wonwoo Shon, Farnaz Dadmanesh, Michele Burnison, Amin Mirhadi, Zachary S. Zumsteg, Rachel Choi, Madison Davis, Joseph Lee, Dustin Rollins, Cynthia Martin, Negin H. Khameneh, Heather McArthur, and Simon R. V. Knott. Single-cell and spatial profiling identify three response trajectories to pembrolizumab and radiation therapy in triple negative breast cancer. 42(1):70–84.e8. ISSN 1535-6108, 1878-3686. doi:10.1016/j.ccell.2023.12.012. URL https://www.cell.com/cancer-cell/abstract/S1535-6108(23)00440-3.

Peter W Harrison, M Ridwan Amode, Olanrewaju Austine-Orimoloye, Andrey G Azov, Matthieu Barba, If Barnes, Arne Becker, Ruth Bennett, Andrew Berry, Jyothish Bhai, Simarpreet Kaur Bhurji, Sanjay Boddu, Paulo R Branco Lins, Lucy Brooks, Shashank Budhanuru Ramaraju, Lahcen I Campbell, Manuel Carbajo Martinez, Mehrnaz Charkhchi, Kapeel Chougule, Alexander Cockburn, Claire Davidson, Nishadi H De Silva, Kamalkumar Dodiya, Sarah Donaldson, Bilal El Houdaigui, Tamara El Naboulsi, Reham Fatima, Carlos Garcia Giron, Thiago Genez, Dionysios Grigo-riadis, Gurpreet S Ghattaoraya, Jose Gonzalez Mar-tinez, Tatiana A Gurbich, Matthew Hardy, Zoe Hollis, Thibaut Hourlier, Toby Hunt, Mike Kay, Vinay Kaykala, Tuan Le, Diana Lemos, Disha Lodha, Diego Marques-Coelho, Gareth Maslen, Gabriela Alejandra Merino, Louisse Paola Mirabueno, Aleena Mushtaq, Syed Nakib Hossain, Denye N Ogeh, Manoj Pandian Sakthivel, Anne Parker, Malcolm Perry, Ivana Piližota, Daniel Poppleton, Irina Prosovetskaia, Shriya Raj, José G Pérez-Silva, Ahamed Imran Abdul Salam, Shradha Saraf, Nuno Saraiva-Agostinho, Dan Shep-pard, Swati Sinha, Botond Sipos, Vasily Sitnik, William Stark, Emily Steed, Marie-Marthe Suner, Likhitha Surapaneni, Kyösti Sutinen, Francesca Floriana Tricomi, David Urbina-Gómez, Andres Veidenberg, Thomas A Walsh, Doreen Ware, Elizabeth Wass, Natalie L Willhoft, Jamie Allen, Jorge Alvarez-Jarreta, Marc Chakiachvili, Bethany Flint, Stefano Giorgetti, Leanne Haggerty, Garth R Ilsley, Jon Keatley, Jane E Loveland, Benjamin Moore, Jonathan M Mudge, Guy Naamati, John Tate, Stephen J Trevanion, Andrea Winterbottom, Adam Frankish, Sarah E Hunt, Fiona Cunningham, Sarah Dyer, Robert D Finn, Fergal J Martin, and Andrew D Yates. Ensembl 2024. 52(D1):D891–D899. ISSN 0305-1048. doi:10.1093/nar/gkad1049. URL https://doi.org/10.1093/nar/gkad1049.

Grace X. Y. Zheng, Jessica M. Terry, Phillip Belgrader, Paul Ryvkin, Zachary W. Bent, Ryan Wilson, Solongo B. Ziraldo, Tobias D. Wheeler, Geoff P. McDermott, Junjie Zhu, Mark T. Gregory, Joe Shuga, Luz Montesclaros, Jason G. Underwood, Donald A. Masquelier, Stefanie Y. Nishimura, Michael Schnall-Levin, Paul W. Wyatt, Christopher M. Hindson, Rajiv Bharadwaj, Alexander Wong, Kevin D. Ness, Lan W. Beppu, H. Joachim Deeg, Christopher McFarland, Keith R. Loeb, William J. Valente, Nolan G. Ericson, Emily A. Stevens, Jerald P. Radich, Tarjei S. Mikkelsen, Benjamin J. Hindson, and Jason H. Bielas. Massively parallel digital transcriptional profiling of single cells. 8(1):14049. ISSN 2041-1723. doi:10.1038/ncomms14049. URL https://www.nature.com/articles/ncomms14049.

Zoe A. Clarke and Gary D. Bader. MALAT1 expression indicates cell quality in single-cell RNA sequencing data. URL https://www.biorxiv.org/content/10.1101/2024.07.14.603469v2.

Tomàs Montserrat-Ayuso and Anna Esteve-Codina. High content of nuclei-free low-quality cells in reference single-cell atlases: A call for more stringent quality control using nuclear fraction. 25(1):1124. ISSN 1471-2164. doi:10.1186/s12864-024-11015-5. URL https://doi.org/10.1186/s12864-024-11015-5.

Michael M. Danziger, Bharath Dandala, Viatcheslav Gurev, Matthew Madgwick, Sivan Ravid, Tim Rumbell, Akira Koseki, Tal Kozlovski, Ching-Huei Tsou, Ella Barkan, Tanwi Biswas, Jielin Xu, Yishai Shimoni, Jianying Hu, and Michal Rosen-Zvi. Bmfm-rna: whole-cell expression decoding improves transcriptomic foundation models, 2026. URL https://arxiv.org/abs/2506.14861.

Tsung-Yi Lin, Priya Goyal, Ross Girshick, Kaiming He, and Piotr Dollár. Focal loss for dense object detection. 42(2):318–327. ISSN 1939-3539. doi:10.1109/TPAMI.2018.2858826. URL https://ieeexplore.ieee.org/document/8417976.

Mukund Sundararajan, Ankur Taly, and Qiqi Yan. Axiomatic attribution for deep networks. Pau Badia-i Mompel, Jesús Vélez Santiago, Jana Braunger, Celina Geiss, Daniel Dimitrov, Sophia Müller-Dott, Petr Taus, Aurelien Dugourd, Christian H Holland, Ricardo O Ramirez Flores, and Julio Saez-Rodriguez. de-coupler: ensemble of computational methods to infer biological activities from omics data. Bioinformatics Advances, 2(1):vbac016. 03 2022. ISSN 2635-0041. doi:10.1093/bioadv/vbac016. URL https://doi.org/10.1093/bioadv/vbac016.

Sophia Müller-Dott, Eirini Tsirvouli, Miguel Vazquez, Ricardo O Ramirez Flores, Pau Badia-i Mompel, Robin Fallegger, Dénes Türei, Astrid Lægreid, and Julio Saez-Rodriguez. Expanding the coverage of regulons from high-confidence prior knowledge for accurate estimation of transcription factor activities. Nucleic Acids Research, 51(20):10934–10949, 10 2023. ISSN 0305-1048. doi:10.1093/nar/gkad841. URL https://doi.org/10.1093/nar/gkad841.

Juanjuan Wang, Ningning Zhu, Xiaomin Su, Yunhuan Gao, and Rongcun Yang. Novel tumor-associated macrophage populations and subpopulations by single cell rna sequencing. Frontiers in immunology, 14: 1264774, 2024.

Yue Xi, Yingchun Zhang, Kun Zheng, Jiawei Zou, Lv Gui, Xin Zou, Liang Chen, Jie Hao, and Yiming Zhang. A chemotherapy response prediction model derived from tumor-promoting b and tregs and proinflammatory macrophages in hgsoc. Frontiers in Oncology, 13: 1171582, 2023.

Thuy Linh Dang Cao, Kunio Kawanishi, Sachie Hashimoto, Kowit Hengphasatporn, Chiaki Nagai-Okatani, Takaharu Kimura, Mohammed Abdelaziz, Rie Shiratani, Thanasis Poullikkas, Nuriza Ulul Azmi, et al. Tumor-expressed gpnmb orchestrates siglec-9+ tam polarization and emt to promote metastasis in triple-negative breast cancer. Proceedings of the National Academy of Sciences, 122(36):e2503081122, 2025.

Kristin Wollenick, Jun Hu, Glen Kristiansen, Peter Schraml, Hubert Rehrauer, Utta Berchner-Pfannschmidt, Joachim Fandrey, Roland H. Wenger, and Daniel P. Stiehl. Synthetic transactivation screening reveals ETV4 as broad coactivator of hypoxia-inducible factor signaling. 40(5):1928–1943. ISSN 0305-1048. doi:10.1093/nar/gkr978. URL https://pmc.ncbi.nlm.nih.gov/articles/PMC3300025/.

Ruben Bill, Pratyaksha Wirapati, Marius Messemaker, Whijae Roh, Beatrice Zitti, Florent Duval, Máté Kiss Jong Chul Park, Talia M. Saal, Jan Hoelzl, David Tarussio, Fabrizio Benedetti, Stéphanie Tissot, Lana Kandalaft, Marco Varrone, Giovanni Ciriello, Thomas A. McKee, Yan Monnier, Maxime Mermod, Emily M. Blaum, Irena Gushterova, Anna L. K. Gonye, Nir Hacohen, Gad Getz, Thorsten R. Mempel, Al-lon M. Klein, Ralph Weissleder, William C. Faquin, Peter M. Sadow, Derrick Lin, Sara I. Pai, Moshe Sade-Feldman, and Mikael J. Pittet. CXCL9:SPP1 macrophage polarity identifies a network of cellular programs that control human cancers. 381(6657):515–524, a. doi:10.1126/science.ade2292. URL https://www.science.org/doi/10.1126/science.ade2292.

Bin-Zhi Qian and Jeffrey W. Pollard. Macrophage Diversity Enhances Tumor Progression and Metastasis. 141(1):39–51. ISSN 0092-8674, 1097-4172. doi:10.1016/j.cell.2010.03.014. URL https://www.cell.com/cell/abstract/S0092-8674(10)00287-4.

Antoni Ribas and Jedd D. Wolchok. Cancer immunotherapy using checkpoint blockade. 359(6382):1350–1355. doi:10.1126/science.aar4060. URL https://www.science.org/doi/10.1126/science.aar4060.

Peng Jiang, Shengqing Gu, Deng Pan, Jingxin Fu, Avinash Sahu, Xihao Hu, Ziyi Li, Nicole Traugh, Xia Bu, Bo Li, Jun Liu, Gordon J. Freeman, Myles A. Brown, Kai W. Wucherpfennig, and X. Shirley Liu. Signatures of T cell dysfunction and exclusion predict cancer immunotherapy response. 24(10):1550–1558. ISSN 1546-170X. doi:10.1038/s41591-018-0136-1. URL https://www.nature.com/articles/s41591-018-0136-1.

Imran G. House, Peter Savas, Junyun Lai, Amanda X.Y. Chen, Amanda J. Oliver, Zhi L. Teo, Kirsten L. Todd, Melissa A. Henderson, Lauren Giuffrida, Emma V. Petley, Kevin Sek, Sherly Mardiana, Tuba N. Gide, Camelia Quek, Richard A. Scolyer, Georgina V. Long, James S. Wilmott, Sherene Loi, Phillip K. Darcy, and Paul A. Beavis. Macrophage-Derived CXCL9 and CXCL10 Are Required for Antitumor Immune Responses Following Immune Checkpoint Blockade. 26 (2):487–504. ISSN 1078-0432. doi:10.1158/1078-0432.CCR-19-1868. URL https://doi.org/10.1158/1078-0432.CCR-19-1868.

Denarda Dangaj, Marine Bruand, Alizée J. Grimm, Catherine Ronet, David Barras, Priyanka A. Duttagupta, Evripidis Lanitis, Jaikumar Duraiswamy, Janos L. Tanyi, Fabian Benencia, Jose Conejo-Garcia, Hena R. Ramay, Kathleen T. Montone, Daniel J. Powell, Phyllis A. Gimotty, Andrea Facciabene, Donald G. Jackson, Jeffrey S. Weber, Scott J. Rodig, Stephen F. Hodi, Lana E. Kandalaft, Melita Irving, Lin Zhang, Periklis Foukas, Sylvie Rusakiewicz, Mauro Delorenzi, and George Coukos. Cooperation between Constitutive and Inducible Chemokines Enables T Cell Engraftment and Immune Attack in Solid Tumors. 35(6):885–900.e10. ISSN 1535-6108, 1878-3686. doi:10.1016/j.ccell.2019.05.004. URL https://www.cell.com/cancer-cell/abstract/S1535-6108(19)30242-9.

Robin Reschke and Thomas F. Gajewski. CXCL9 and CXCL10 bring the heat to tumors. 7(73):eabq6509. doi:10.1126/sciimmunol.abq6509. URL https://www.science.org/doi/abs/10.1126/sciimmunol.abq6509.

Paola Marie Marcovecchio, Graham Thomas, and Shahram Salek-Ardakani. CXCL9-expressing tumor-associated macrophages: New players in the fight against cancer. 9(2). ISSN 2051-1426. doi:10.1136/jitc-2020-002045. URL https://jitc.bmj.com/content/9/2/e002045.

Ruben Bill, Pratyaksha Wirapati, Marius Messemaker, Whijae Roh, Beatrice Zitti, Florent Duval, Máté Kiss Jong Chul Park, Talia M. Saal, Jan Hoelzl, David Tarussio, Fabrizio Benedetti, Stéphanie Tissot, Lana Kandalaft, Marco Varrone, Giovanni Ciriello, Thomas A. McKee, Yan Monnier, Maxime Mermod, Emily M. Blaum, Irena Gushterova, Anna L. K. Gonye, Nir Hacohen, Gad Getz, Thorsten R. Mempel, Al-lon M. Klein, Ralph Weissleder, William C. Faquin, Peter M. Sadow, Derrick Lin, Sara I. Pai, Moshe Sade-Feldman, and Mikael J. Pittet. CXCL9:SPP1 macrophage polarity identifies a network of cellular programs that control human cancers. 381(6657):515–524, a.doi:10.1126/science.ade2292. URL https://www.science.org/doi/10.1126/science.ade2292.

Ryuma Tokunaga, Wu Zhang, Madiha Naseem, Alberto Puccini, Martin D. Berger, Shivani Soni, Michelle McSkane, Hideo Baba, and Heinz-Josef Lenz. CXCL9, CXCL10, CXCL11/CXCR3 axis for immune activation – A target for novel cancer therapy. 63:40–47. ISSN 0305-7372, 1532-1967. doi:10.1016/j.ctrv.2017.11.007. URL https://www.cancertreatmentreviews.com/article/S0305-7372(17)30199-8/abstract.

Qiang Ding, Panpan Lu, Yujia Xia, Shuping Ding, Yuhui Fan, Xin Li, Ping Han, Jingmei Liu, Dean Tian, and Mei Liu. CXCL9: Evidence and contradictions for its role in tumor progression. 5(11):3246–3259. ISSN 2045-7634. doi:10.1002/cam4.934. URL https://onlinelibrary.wiley.com/doi/abs/10.1002/cam4.934.

Mahnoor Gondal, Marcin Cieslik, and Arul Chinnaiyan. Integrated cancer cell-specific single-cell RNA-seq datasets of immune checkpoint blockade-treated patients, b. URL https://zenodo.org/records/14511579.

